# Polygenic backgrounds influence phenotypic consequences of variants in cells, individuals, and populations

**DOI:** 10.1101/2025.01.07.631805

**Authors:** Madison Chapel, Jessica Dennis, Carl G. de Boer

## Abstract

Both rare and common genetic variants contribute to human disease, and emerging evidence suggests that they combine additively to influence disease liability. However, the non-linear relationship between disease liability and disease prevalence means that risk variants may have more severe phenotypic consequences in high-risk polygenic backgrounds and minimal impact in low-risk backgrounds, resulting in uneven selection across the population. As a result, selection coefficients may be better modeled as distributions that differ across populations, time, environments, and individuals rather than single values. Further, the number of genes contributing to a trait and epistasis between alleles enhance negative selection due to the increased variance pushing more individuals to phenotypic extremes. Because disease-relevant phenotypes may be masked in certain genetic backgrounds, polygenic background should be considered when characterizing the molecular underpinnings of complex traits.

## Introduction

Many genetic diseases are influenced by both rare and common variants. Highly penetrant variants often follow Mendelian inheritance patterns. These variants are typically rare, occurring at low allele frequencies throughout the population^1,2^. Many other diseases are polygenic, with common genetic variants of small effect combining to influence disease risk, resulting in a spectrum of genetic risk across a population^3^. Genome-Wide Association Studies (GWAS) have become crucial for understanding how these common variants influence disease risk^4,5^. While rare and common variants are often discussed as though they are distinctly different categories, as we discuss below, both types of variants fall along opposite ends of a shared effect size spectrum.

By aggregating effect estimates from GWAS summary statistics, researchers can calculate Polygenic Scores (PGS), also called Polygenic Risk Scores (PRS), estimating an individual’s genetic liability for a trait or disease based on their genotype^6,7^. For binary traits, like diagnosis of a disease, PGSs are typically calculated as the sum of alleles, with each allele weighted by the log odds ratio of developing disease when that allele is present. Even with increasingly large cohort sizes, GWAS have limited power to detect effects from rare variants, and so PGSs typically capture only the influence of common genetic variation^5,8^. Similarly, rare variants are often studied in isolation, without consideration of the genetic background in which they occur. Some potentially causal variants are inherited without consistently leading to disease or occur at frequencies incompatible with causing a highly penetrant Mendelian disease^9^. Moreover, individuals with the same causal alleles may show variable expressivity of the disease phenotype^10^.

Several recent studies have characterized cohorts for both rare disease variants and PGS for various genetic diseases, including breast and ovarian cancer^11,12^, developmental disorders^13^, Parkinson’s disease^14^, and others^15–18^. The combination of high PGS and rare variant results in a higher disease risk than either factor alone^12,17^. When examining the joint effect of rare and common genetic variants, several studies demonstrated that these two sources of genetic variation combine roughly additively to determine the log odds ratios of disease^11,13–16^, essentially meaning that the rare variant effects are additive in the same way as the common alleles of a PGS (albeit with a relatively large weight). This additive trend was robust across multiple studies, despite differences in cohort size and approaches to PGS stratification. Importantly, while an additive PGS could still be predictive even if the alleles combine non-additively to impact disease (e.g. by capturing their average effects), the large effect sizes of the rare variants in these studies provide a much more robust test of the additive model, and the rare variants’ effects on log odds of disease were similar regardless of polygenic background.

These studies also highlight a key concept: while variant effects may be additive, their phenotypic consequences can vary depending on the genetic background in which they occur. Disease prevalence is often modeled as a function of underlying liability, which reflects the combined influences of genetic and environmental factors^19–21^. While the mapping between prevalence and liability is non-linear, it can be approximated linearly through transformation to log odds ratios^22^, where variant effects are additive. Within this framework, the penetrance of a rare variant may vary across polygenic backgrounds. In a low-risk background, a variant may seldom lead to disease, while in a high-risk background, the same variant may push liability past a disease threshold. This background dependence can influence not only disease prevalence, but also age of onset and disease severity^14,16,23^.

In this paper, we explore the consequences of how variants combine to influence selective pressures, and present key considerations for researchers investigating variant effects in cellular models, where polygenic background is often unaccounted for.

### Penetrance and expressivity of rare variants can be modified by genetic background

Although genetic risk factors combine additively on polygenic scores and disease liability, this does not imply additivity on disease prevalence or probability (**Fig. 1; Methods**). As described above, genetic studies have demonstrated that disease variants combine additively on the log odds ratios scale, meaning that they act multiplicatively on odds ratios^24^. Consequently, a variant is predicted to have the same effect on the log odds ratio of disease regardless of the genetic background it arises in but have very different effects on disease prevalence, where selection is acting.

**Figure 1.**
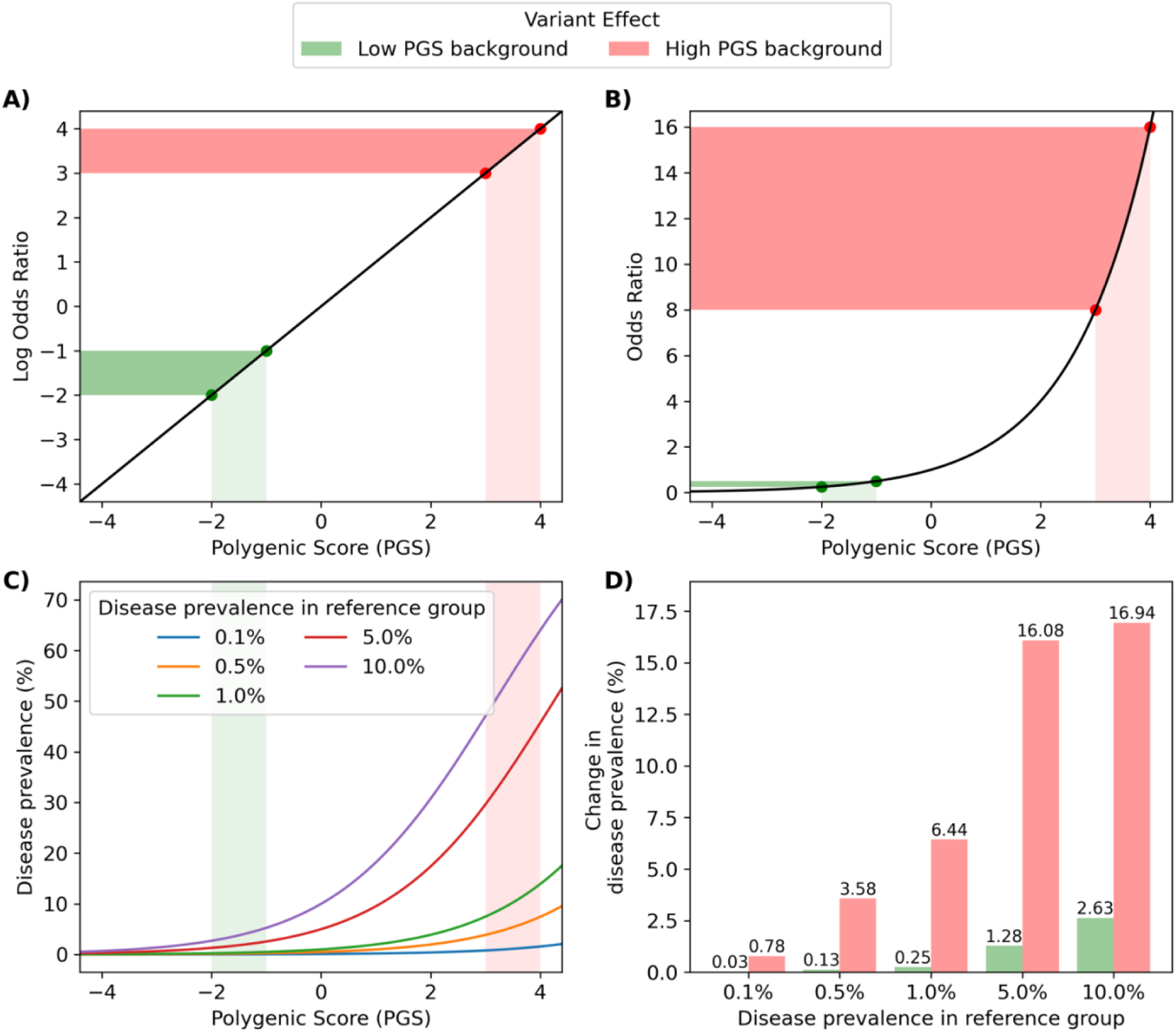
Variant effects depend on polygenic background. (**A**) We simulated polygenic scores (PGS, x axis) that are linearly related to log odds ratios of disease (y axis), resulting in a risk variant having the same change in log odds ratio regardless of where on the PGS distribution the individual lies (colors). (**B**) However, similar changes to PGS (x axis) result in very different changes to odds ratios (y axis) depending on where on the PGS distribution the individual lies (colors). **(C)** The relationship between PGS (x axis) and disease prevalence (y axis) is also influenced by the background rate of disease in the population (colors), with populations already at higher risk of disease experiencing greater increases at the extreme end of the PGS distribution. **(D)** Disease prevalence (y axis) increases substantially more when risk variants are introduced to high PGS backgrounds (red bars) than low PGS backgrounds (green bars), regardless of the disease background rate (x axis).

To demonstrate how variants’ impacts on fitness depends on polygenic background, we created a simulation of genetic disease. Here, PGSs are linearly proportional to log odds ratios, and disease prevalence is modeled by multiplying this odds ratio by the odds of disease in the reference group and transforming back to a probability (**Methods**). In this framework, a large-effect variant that increases the PGS by one unit increases the log odds ratio of disease by the same amount regardless of genetic background (**Fig. 1A**). That same variant, however, will result in a minimal change in odds ratio in a low-risk background and a dramatic change in a high-risk background (**Fig. 1B**). Consequently, the changes in absolute disease prevalence associated with the variant vary widely depending on the genetic background in which it occurs. (**Fig. 1C,D**).

### Genetic background modifies the strength of selection

Selection for or against a genetic variant is dependent on the variant’s impact on fitness, which is in turn mediated by its effect on disease. However, this relationship is not uniform across all traits or genetic backgrounds. As demonstrated above (**Fig. 1**), variants are expected to have very different impacts on disease risk depending on polygenic background. Consequently, the same variant presenting in different genetic backgrounds can also be expected to have different fitness consequences.

Not all diseases impact fitness in the same way. In cases like immunity, there is a balance between immunity against pathogens and autoimmunity, and either extreme can be disfavored^24–26^. In cases where disease diagnosis depends on a quantitative trait surpassing a threshold, such as high blood pressure^27^ or obesity^28^, there may be a more proportional relationship between disease liability and selection^29,30^. In cases where the disease is characterized by a catastrophic event, like cancer diagnosis or sudden cardiac arrest, there may be some relationship between disease liability and fitness before the catastrophic event, but the event itself, often resulting in death or impairment, clearly has a direct and substantial impact on survival. The mapping from liability to fitness can differ depending on the trait, and selection will subsequently act more strongly in some contexts than others. Regardless of disease, the relationship between a liability with additive genetic contributions and fitness must be non-linear because fitness is bounded between 0 and 1 (unable to reproduce and maximal reproduction, respectively).

When a population inherently has a low genetic liability of disease, a pathogenic variant – even one of large effect – may rarely cause disease. Selection against the pathogenic variant may be driven primarily by individuals at the high end of the risk distribution, where the probability of disease is highest. In low-risk individuals, the variant may have minimal phenotypic consequences and so remain effectively neutral in these individuals, reducing the ability of selection to eliminate the allele^31,32^. Selection pressures can also change over time due to shifts in allele frequencies or environmental conditions. Even in controlled experiments, cryptic environmental variation can cause selection coefficient estimates to differ between replicates^33^. Accordingly, Gallet et al proposed representing selection coefficients as a distribution instead of a single value^33^. Similarly, selection coefficients for natural populations may also be better captured as a distribution that changes to account for the population’s allele frequencies and environmental conditions.

Intriguingly, the strength of selection is increased as more genes contribute to the trait, due to increased variance in the genetic liability distribution. The number of genes contributing to a trait can change over time as the molecular underpinnings evolve, for instance, via transcription factor binding site gain, increasing gene regulatory network connectivity and producing new conduits through which variants can impact the trait^34^. We simulated this by testing various numbers of genes contributing to a trait, where each gene had alleles whose frequencies and effect sizes were drawn from the same underlying distribution (**Methods**). While the means of the distributions remained unchanged, the populations with more genes contributing to a trait have more individuals with extreme polygenic scores (**Fig. 2A, B**), and thus extreme phenotypes, due to the increased variance in the distribution. The variance increased linearly with the number of genes contributing to the trait (**Fig. 2B**), consistent with a sum of independent binomial random variables (probabilities are allele frequencies, and, in our case, weights are effect sizes).

**Figure 2:**
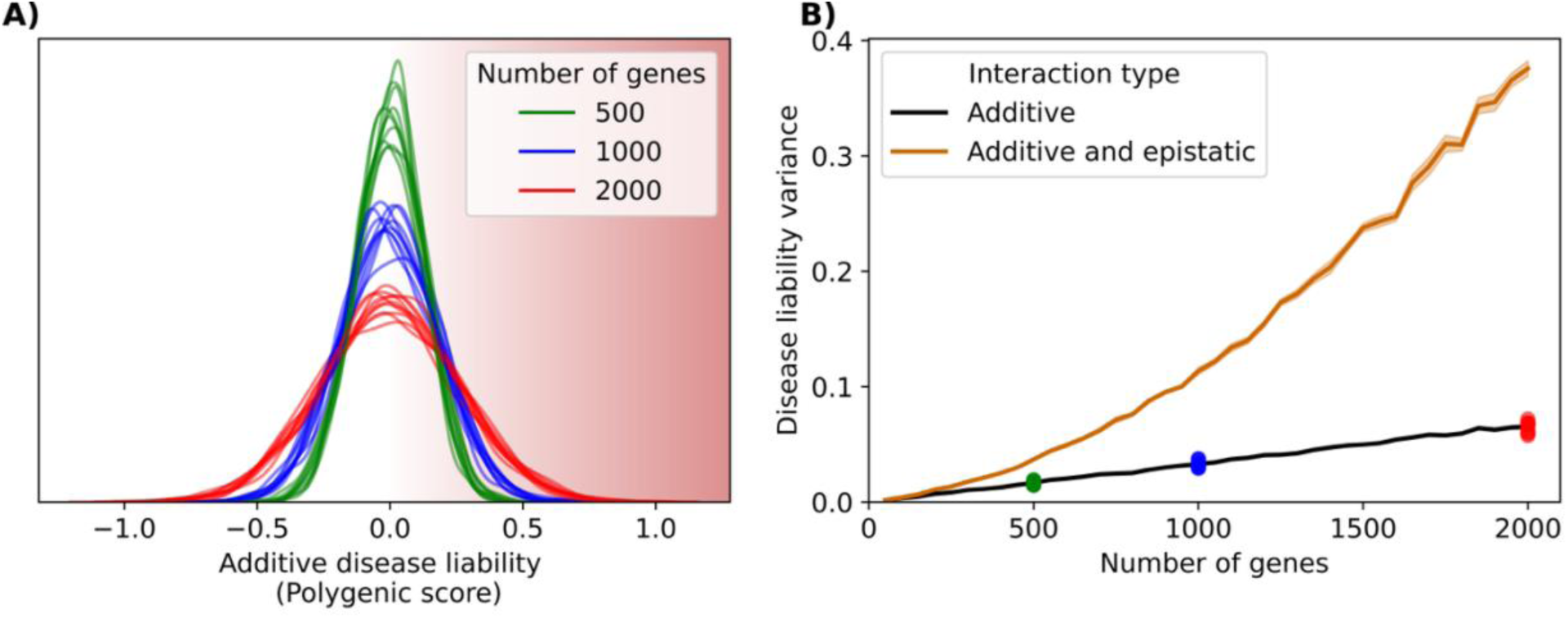
Variation in disease risk increases linearly with the number of contributing genes for additive traits and super-linearly with epistasis. (**A**) Density of individuals (y axis) of disease risk polygenic scores (x axis) for individuals in simulated populations that are matched for population size, effect size distributions, and allele frequencies, but differ by the number of contributing genes (colors). Individuals at extremes are more likely to develop disease (red shading). (**B**) PGS variance (y axis) for each number of genes (x axis) for both simulating strictly additive interactions (black) and including epistatic interactions (orange). Lines represent means across 10 replicates, shaded regions show the standard error of the mean. Colored dots are the additive PGS variance values for the correspondingly colored populations in (**A**).

In contrast to this simulation, populations are continuously evolving. Selection will continually act against individuals at the high-risk end of the distribution. Since large-effect variants are over-represented at this tail^35^, the allele frequencies of high-risk variants will decrease, which will result in more phenotypic variation being explained by low-risk variants. Consequently, negative selection drives effect sizes of alleles in the population to be increasingly small and homogenous for traits with many contributing genes^20,36^. Interestingly, this “flattening” induced by negative selection^36^, increases the apparent polygenicity of a trait because more of the trait’s heritability is explained by a greater number of variants, but we show that increasing the polygenicity in terms of the molecular architecture underpinning the trait increases the power of negative selection by pushing more individuals to trait extremes.

This simulation also assumes additive contributions to disease liability from each variant as this is most consistent with the clinical genotype-phenotype data (reviewed above). However, if variants were to interact super-additively on genetic liability (e.g. multiplicatively), the variance in the disease liability distribution would be far greater, pushing more individuals to phenotypic extremes (**Fig. 2B**; **Methods**). Such combinations may be subject to strong negative selection, reducing the frequency or effect sizes of interacting alleles. This may contribute to the apparent difficulty in detecting epistatic interactions in natural populations^37,38^, despite their prevalence in synthetic systems^39,40^.

The environment a population occupies can modify the way that selection acts on genetic variation and can be thought of as an independent modifier of disease liability. If a population moves to an environment that predisposes them to disease, the liability distribution will shift, increasing the number of individuals at high risk of developing disease. More individuals in the population would be subject to negative selection, reducing the frequency of disease-associated variants. Termed “polygenic adaptation”, allele frequencies continue to change over time to approach the new fitness optimum, leading to a population with reduced genetic liability^41,42^. If environmental pressures relax again, the overall liability distribution shifts back, but the population now has a lower disease liability than at the outset due to the change in allele frequencies (**Figure 3**). A potential example of this has been observed in Inuit populations. Inuit populations appear to have been under selection at the *FADS* region, which encodes several genes involved in fatty acid metabolism^43,44^, potentially reflecting adaptation to their traditional diet high in polyunsaturated fatty acids. In European populations, variants in *FADS1* are associated with lower blood cholesterol, suggesting a potential protective effect against cardiometabolic phenotypes^43^. Although, whether the *FADS* variants enriched in Inuit populations translate to lower incidence of cardiometabolic phenotypes, particularly in populations adopting a more Westernized diet with reduced fatty acid content (i.e., a shift in environmental risk factors), remains to be tested^45^.

**Figure 3:**
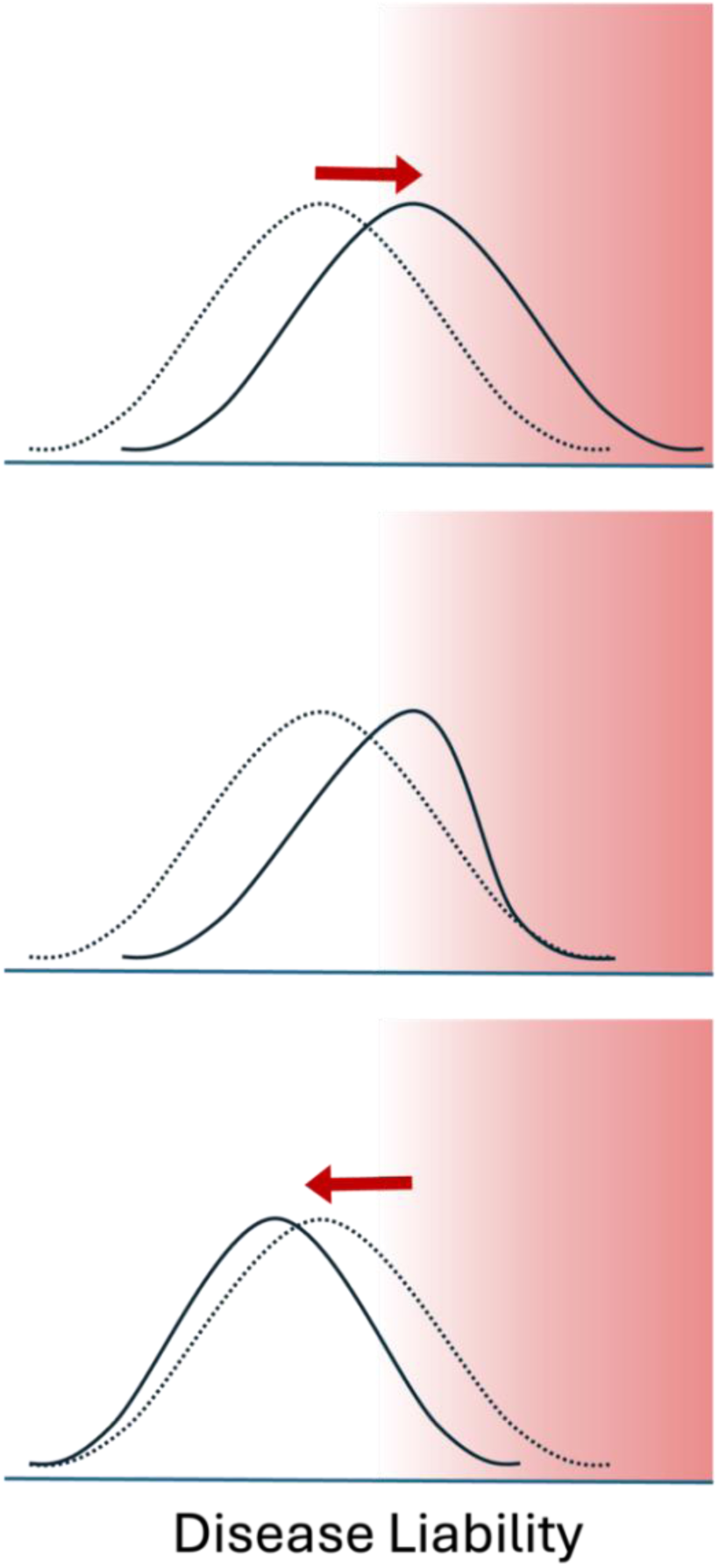
The distribution of disease liability in a population can change in response to environmental shifts. A change in the environment (top panel) shifts the original distribution (dashed line) to higher disease liability (solid line), predisposing more individuals to disease (red shading). In the new environment, more individuals have a high probability of developing disease, and selection will eliminate those at highest risk (middle panel), changing the disease liability distribution within a population. The environmental stressor is removed (bottom panel), resulting in a population (solid line) with lower mean disease liability than the original (dotted line).

Many drugs act as specific modifiers of gene function and can also act as an independent modifier of disease liability. For instance, cancer patients undergoing immune checkpoint blockade, where immune-dampening pathways (e.g. PD1/PD-L1 CTLA4) are inhibited, are more likely to experience immune-related adverse events if they are also genetically predisposed to autoimmunity (e.g. via a high autoimmunity PGS)^46–48^. In another example, tyrosine kinase inhibitors – typically used as anticancer agents – are more toxic to cardiomyocytes derived from Long QT syndrome patients than those derived from healthy controls. This also suggests that drugs or other environmental stressors could be used to draw out variant effects. Phenotypes associated with risk variants that may be masked in one environment could be revealed when an environmental stressor is applied. If the environment influences disease liability, it will also modify the strength of selection and the allele frequency distribution.

### Implications for functional studies investigating the genetic basis of complex traits

Characterizing the mechanisms underlying GWAS variants using functional assays is now a major focus of human genetics research^51^. Obviously, using the correct cell type for the assay is critical to determining the true mechanisms underlying a variant’s role in disease. Using an appropriate genetic background may also prove to be crucial in many circumstances as the variant’s phenotypic effect could be masked by the genetic background. If a cell line used for functional characterization is derived from a low-polygenic risk background, certain variants may have no observable effect when looking at broad measures such as cellular phenotypes or even gene expression (as in shadow enhancer genes; see below). Were the same variant to be studied in a cell line derived from a higher-risk background, relevant changes in phenotype may become apparent. This mirrors observations from a population level, where the same rare variant can result in variable impacts on disease risk depending on the individual’s genetic background. For instance, variation elsewhere in the genome may explain why cardiomyocytes derived from symptomatic and asymptomatic sibling carriers of the same Long QT syndrome-associated variant displayed varying responses to QT interval-modifying drugs^52,53^.

Functional studies may be improved by considering the cellular model’s PGSs for relevant traits and perhaps replicating results in alternate cellular models with diverse PGSs. This would allow researchers to investigate not only whether variant effects are revealed in high-risk backgrounds, but also how genetic background modulates the expressivity of a variant. A key question is whether the phenotypic change associated with a variant is constant across backgrounds, or whether it increases more drastically in higher-risk contexts, mirroring the relationship between disease liability and disease risk at the population level (**Fig. 1)**. Similarly, a variant may result in a step-like function, with little change in phenotype when polygenic risk is sufficiently low but a large change after a certain threshold, suggesting that the phenotypic changes at the cellular level act according to a liability-threshold model^19^. Performing assays across a range of polygenic backgrounds could help to distinguish these models.

Cell lines with high polygenic scores for the trait of interest will likely be the most informative, as they will maximize the phenotypic effects of perturbations. However, strongly deleterious alleles may be better studied in low-risk polygenic backgrounds where phenotypes may be less extreme, facilitating more complete characterization (e.g. the risk allele and high-PGS combination is not viable). This is supported by clinical studies, where rare variants of large effect are more common in patients with lower PGSs^15,54,55^. With the growing use of patient-derived induced pluripotent stem cells (iPSCs) as a model to research genetic disease^56^, iPSC banks such as iPSCORE, which include not only phenotype data but also matched genotyping data, may prove invaluable for future research endeavors^57^. Including pre-calculated PGSs will further increase the utility of these iPSC banks by enabling researchers to quickly identify the lines most likely to reveal variant effects.

Shadow enhancers provide an intriguing example of how genetic background can sensitize mutation effects. Shadow, or distributed enhancers, are sets of enhancers that regulate a common target and drive overlapping expression patterns, seemingly with redundant functions^58^. However, this redundancy may simply reflect the genetic background of the strain being tested. Most typically, such experiments are done in inbred lines, and whether the enhancers appear to be redundant may depend on whatever polygenic background happens to have been fixed. For instance, in a natural population, a mutation disrupting the activity of one of a pair of shadow enhancers may not be tolerated if genetic variation elsewhere in the genome (e.g. in transcriptional regulators) disrupts the activity of the remaining enhancer, resulting in a fitness defect. Indeed, deleting three of six enhancers for the *shavenbaby* gene had no effect on *D. melanogaster* development under normal conditions but caused phenotypic changes in mutants with a single copy of the *wingless* gene^59^. Similar results were shown in enhancers regulating *Pax6*^60^, *Shh*^61^, and several loci involved in limb development^62^. Similar circumstances could mask the effects of GWAS variants, the vast majority of which are non-coding and presumed to be regulatory^63,64^, and many of which affect the expression of transcriptional regulators^35,65^.

## Conclusions

The relationship between variants and disease risk can be modified by such factors as an individual’s age, environment, or sex chromosome composition, and there is a growing initiative to study and characterize these interactions^18,66,67^. The context of polygenic background is similarly important but seldom discussed. As we show, polygenic background can sensitize cells, individuals, or populations to variant effects.

The dependence on polygenic background necessitates a shift in how we discuss and study disease-associated variants. Instead of associating each variant with a discrete effect on disease risk, each variant should instead be conceptualized as having a context-dependent distribution of effects. Some recent predictive tools have already begun to reflect this shift: for example, RICE^68^ combines rare and common variant information into a single genetic liability predictor, and has demonstrated more accurate predictions of risk than traditional PRS-only approaches.

Recognizing that variant effects are context-dependent is critical not only for building more realistic models of genetic risk, but also for how we validate variant effects experimentally. Functional assays, genetic association studies, and clinical interpretation efforts must increasingly account for the fact that phenotype consequences may differ across genetic backgrounds. Incorporating polygenic background will be necessary to understand the evolutionary origins and causal mechanisms underlying complex traits.

## Methods

### Modeling how polygenic background can modify changes in disease risk associated with rare variants

To demonstrate how the effect of a variant changes depending on the genetic background in which it occurs (**Fig. 1**), a range of polygenic scores between –4 and 4 were plotted in log_2_ space (log odds ratio; log(OR)) and linear space (odds ratio; OR). A large-effect variant with an effect size of one PGS unit was modeled by shifting the PGS from -2 to -1 for a low-risk background, and from 3 to 4 for a high-risk background (**Fig. 1A,B**).

Disease prevalence across the range of PGS (**Fig. 1C,D**) was calculated by first determining the odds of disease in the reference group based on disease prevalence *(p*) for five different background rates of disease:

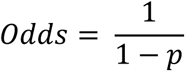

We then determined disease prevalence across the range of corresponding odds ratios (*OR*):

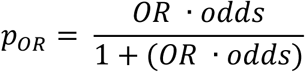

### Impact of polygenicity degree on polygenic score distributions

To model how variance in the PGS distribution increases as more genes contribute to the disease trait (**Fig. 2**), sample populations were generated for increasing numbers of genes, ranging from 50 to 2000 in increments of 50. For each gene count, 10 replicate populations of 1000 individuals were generated. Allele frequencies for each replicate were generated from a uniform distribution *U*(0.0, 1.0). Additive effect sizes, *β*, were generated from a normal distribution *N*(mean = 0.0, standard deviation = 0.01). Epistatic effect sizes, *γ*, which are expected to be much smaller than additive effects^69^, were generated from a normal distribution *N*(mean = 0.0, standard deviation = 0.0004). Individual genotypes, *χ*, were simulated using a binomial distribution with *n*=2 and *p* equal to the generated allele frequencies.

In the additive model, disease liability, *L*, was determined by summing the products of effect sizes and genotypes:

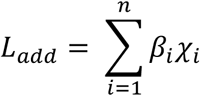

In model including epistatic interactions, liability was calculated as:

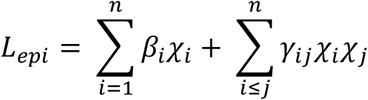

Where *γ_ij_* is the interaction effect between SNPs *i* and *j*. This model includes only additive effects and additive × additive (i.e., epistatic) interaction effects but omits dominance-related interaction terms^70,71^.

## Acknowledgements

We thank S. Otto and Z. Laksman for helpful discussions. This research was supported by the Natural Sciences and Engineering Research Council of Canada (RGPIN-2020-05425), and the Canadian Institute for Health Research (PJT-180537). M.C. was supported by a UBC IGF. C.G.D. is a Michael Smith Health Research BC Scholar.

## Declaration of interests

The authors declare no competing interests.

## Notes

### Competing Interest Statement

The authors have declared no competing interest.

### Summary of Updates

Substantial modifications have been made throughout the manuscript to improve the presentation of our results and better contextualize them within the current context of the field. Sections of the manuscript have been rearranged so that it concludes with the most forward-looking content. Additional detail has been added to the text to better explain the disease liability simulations, and new data for disease liability with an epistatic interaction model has been added. Additional references have been added to better support assumptions about variant effect additivity in different disease contexts and how disease liability interacts with selection.

